# SUMOnet: Deep Sequential Prediction of SUMOylation Sites

**DOI:** 10.1101/2023.08.25.554749

**Authors:** Berke Dilekoglu, Oznur Tastan

## Abstract

SUMOylation is a reversible post-translational protein modification in which SUMOs (small ubiquitin-like modifiers) covalently attach to a specific lysine residue of the target protein. This process is vital for many cellular events. Aberrant SUMOylation is associated with several diseases, including Alzheimer’s, cancer, and diabetes. Therefore, accurate identification of SUMOylation sites is essential to understanding cellular processes and pathologies that arise with their disruption. We present three deep neural architectures, SUMOnets, that take the peptide sequence centered on the candidate SUMOylation site as input and predict whether the lysine could be SUMOylated. Each of these models, SUMOnet-1, -2, and -3 relies on different compositions of deep sequential learning architectural units, such as bidirectional Gated Recurrent Units(biGRUs) and convolutional layers. We evaluate these models on the benchmark dataset with three different input peptide representations of the input sequence. SUMOnet-3 achieves 75.8% AUPR and 87% AUC scores, corresponding to approximately 5% improvement over the closest state-of-the-art SUMOylation predictor and 16% improvement over GPS-SUMO, the most widely adopted tool. We also evaluate models on a challenging subset of the test data formed based on the absence and presence of known SUMOylation motifs. Even though the performances of all methods degrade in these cases, SUMOnet-3 remains the best predictor in these challenging cases.

**Availability and Implementation:** The SUMOnet-3 framework is available as an open-source project and a Python library at https://github.com/berkedilekoglu/SUMOnet.

## 1 Introduction

Most proteins undergo post-translational modifications (PTMs) throughout their lifetimes [46]. PTMs modulate the modified proteins’ functions by altering their targets’ structure, dynamics, subcellular locations, and/or interactions, yielding a greater functional repertoire of the proteome [46]. SUMOylation is one of the most critical post-translational modifications (PTM) in eukaryotic cells, in which small ubiquitin-like modifiers (SUMOs) covalently attach to specific lysine (K) residues of target proteins in a reversible manner [4]. SUMO proteins are ubiquitously expressed in eukaryotes and are highly conserved, indicating their functional importance [16]. SUMO was initially characterized for its role in binding nuclear proteins [34,33]; later, its wide range of activities were discovered in transcription regulation, chromatin remodeling, DNA repair, and the control of cell cycle progression [26,14,4]. Due to its critical role in the regulation of cell cycle and cellular responses to stress conditions, alterations in SUMOylation process have also been associated with several diseases, including Alzheimer’s [30], cancer [41], and autoimmune diseases [51].

SUMO covalently attaches to its target through a series of reactions facilitated by SUMO-activating enzymes 1 (E1), SUMO-conjugating enzyme E2 (Ubc9), and various SUMO E3 ligases. SUMO-specific proteases cleave the bond between SUMO and its substrate to reverse the SUMOylation process [16]. Mass spectrometry (MS)-based proteomics can detect the SUMOylated proteins [45,13,18] and SUMOylation sites [43,21,22] in a high-throughput fashion. However, the transient nature of SUMOylation, the low stoichiometry of SUMO, and the small fraction of the SUMOylated protein pose technical challenges to the probing of SUMOylation events [22]. Therefore, computational methods have been proposed to assist experimental efforts by narrowing down possible SUMOylation sites.

Several SUMOylation site predictors rely on the known SUMOylation motifs or alignments of experimentally validated SUMOylation sites and calculate a score based on these motif presences for prediction. One method that relies on these motifs is JASSA [2]. JASSA groups experimentally validated SUMOylation sites in clusters based on the presence of the consensus motif and its inverted form. It then predicts SUMOylation sites using a scoring system based on the position-specific frequencies of amino acids calculated specifically for each of these groups. There is a fundamental limitation of these motif-based methods: not every sequence that contains the consensus motifs is necessarily SUMOylated, and not every SUMOylated site contains the motif. A second limitation for these is that the position-specific matrices fail to capture interactions across positions. A second widely used tool is GPS-SUMO [52], which is the latest version of a series of tools developed earlier (SUMOsp [49], and SUMOsp 2.0 [40]). GPS-SUMO was developed using the group-based phosphorylation scoring algorithm. Another tool is pSumo-CD [27] and it uses a covariance discriminant algorithm [9] for prediction. SUMO-Forest [38] uses bi-gram and k-skip-bi-gram (with k=1,2) to represent the input peptide. It relies on an ensemble technique, Cascade Forest [53]. C-iSUMO [32] uses an Adaboost classifier [15] that relies on features derived from structural properties such as the accessible surface area of protein site and backbone torsion angles between residues. There are several other methods that rely on various machine learning models [48,47,20,6].

In this work, we introduce three deep sequential learning architectures, SUMOnet-1, -2, and -3, for SUMOylation prediction. These architectures utilize fundamental sequential data processing units such as convolutional layers, GRUs, and LSTMs. We couple these architectures with three different peptide encodings. We compare the proposed SUMOnets against other machine learning methods, such as gradient-boosted trees or logistic regression and other widely adapted SUMOylation predictors on the same benchmark dataset. SUMOnets outperform the existing methods in terms of prediction performance. The performance difference is especially pronounced in challenging cases. We provide the best-performing architecture, SUMOnet-3, as a Python library that can be easily installed.

## 2 Methods

We cast the SUMOylation prediction as a binary classification task in which the input is the 21-mer peptide sequence, *x*, centered on a lysine residue and the class label is given as *y* ∈ − 1, 1. Here, label 1 indicates the positive class where the site is SUMOylated, and -1 indicates the negative class. We propose three alternative architectures, SUMOnet-1, -2, and -3, and train these models in a supervised learning setup. We evaluate the performance of these models with three different peptide input representations.

Below, we first describe the dataset that we use in our experiments. Secondly, we describe the encoding methods. In the subsequent steps, we present the components of the SUMO-nets and the design process of our deep learning architectures SUMOnet-1, -2, and -3, evaluation metrics. Finally, we perform hyperparameter tuning on each model.

### 2.1 Dataset

We obtained the experimentally identified SUMOylation samples from the dbPTM database [25]. We chose this data source for two reasons. Firstly, it is comprehensive as it culls data from various biological databases. Secondly, it provides an up-to-date non-homologous benchmark dataset. Although most prediction tools provide predictive performance results, they use different datasets, most of which contain redundant sequences. Using this benchmark data, we hope others will also be able to compare our methods with theirs. We use the non-redundant benchmark dataset (version 2, downloaded on 12.01.2021). dbPTM database curates this data with the CD-hit program [31] such that no sequence pair have similarity more than 40% [25]. This ensures that the test data do not include sequences very similar to sequences that the model has trained on and to avoid optimistic test performance estimates.

The peptide sequences with the SUMOylated sites constitute the positive set in our classification task. The 21-mer peptides that are also centered on a lysine residue but are not reported to be a SUMOylation site constitute the negative examples. The dataset contains 1,432 proteins and 5,191 SUMOylation positive examples, and 16,066 negative examples in total. This corresponds to a 1:3 ratio of positive-to-negative class labels. We randomly held out 10% of the examples using stratified sampling for class labels and used it as test data to evaluate the models. We tuned models’ hyperparameters on the training data using 5-fold cross-validation. To further evaluate the models, we subset for the most challenging examples in the test data. For this, we subset peptides that contain a SUMOylation motif but are not SUMOylated; these inputs constitute the hard negative SUMOylation cases. We also subset peptide sequences that do not contain a known SUMOylation motif but are SUMOylated. We also provide evaluation results on this hard subset.

### 2.2 Peptide Encodings

To numerically represent the input peptide sequence, we use three different encoding techniques: one-hot, BLOSUM62 [23], and NLF [36]. Each 21-mer peptide can be represented with 21xN vectors. The dimension *N* of the vectors is dependent on the encoding mechanism. They are set as follows:

– **One-hot encoding:** Each candidate SUMOylation site is represented with a 21-mer peptide sequence centered on the lysine residue; hence, the input sequence is represented by a 21 * 21 = 441 dimensional vector.
– **BLOSUM62 encoding**: The BLOSUM62 value for a particular pair of amino acids is the log-odds ratio that estimates the biological probability of substitution to occur relative to that substitution being merely by chance. The row for a particular amino acid is used as an encoding. *N* = 24; thus, each peptide is represented by a 21 * 24 = 504 dimensional vector.
– **NLF encoding**: NLF encoding represents the physicochemical properties of amino acids [36]. The factor is determined using a non-linear Fisher’s transform [36]. A vector of length 18 represents each amino acid. Hence, an input is represented with a 21 * 18 = 378 dimensional vector. Epitopepredict tool developed by [12] is used for extracting NLF representation.

Vector representations of lysine ‘K ‘amino acid in each different encoding are illustrated in Figure 1. We treated the encoding scheme as a hyperparameter and separately selected the best-performing one for each SUMOnet architecture.

**Fig. 1:**
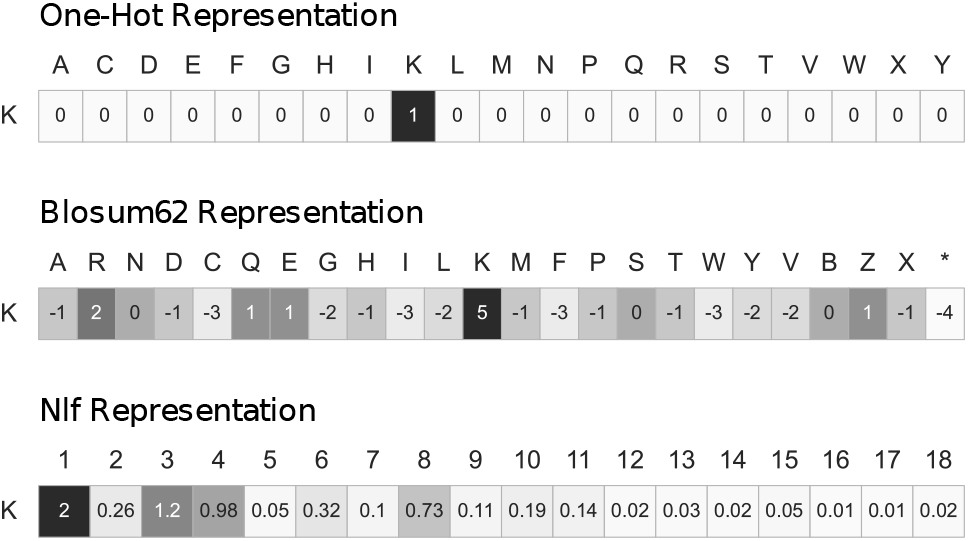
The three encodings used in the paper are shown for the lysine ‘K’ amino acid vector.

### 2.3 SUMOnet Architectures

We train novel deep-learning architectures to classify SUMOylated and non-SUMOylated input peptide sequences. Each SUMOnet architecture is coupled with the best-performing encoding scheme for that specific architecture, which is decided during the hyperparameter selection process. Our architectures consist of combinations of convolutional neural network (CNN) layers [29], Gated Recurrent Units (GRU) [7], bidirectional Gated Recurrent Units (biGRU) [19], self-attention mechanism [44] and feed-forward neural network (FFNN) [37]. We use Keras [8] to implement the architectures. The three SUMO-net architectures are visualized in Figure 2, and we provide details below:

**Fig. 2:**
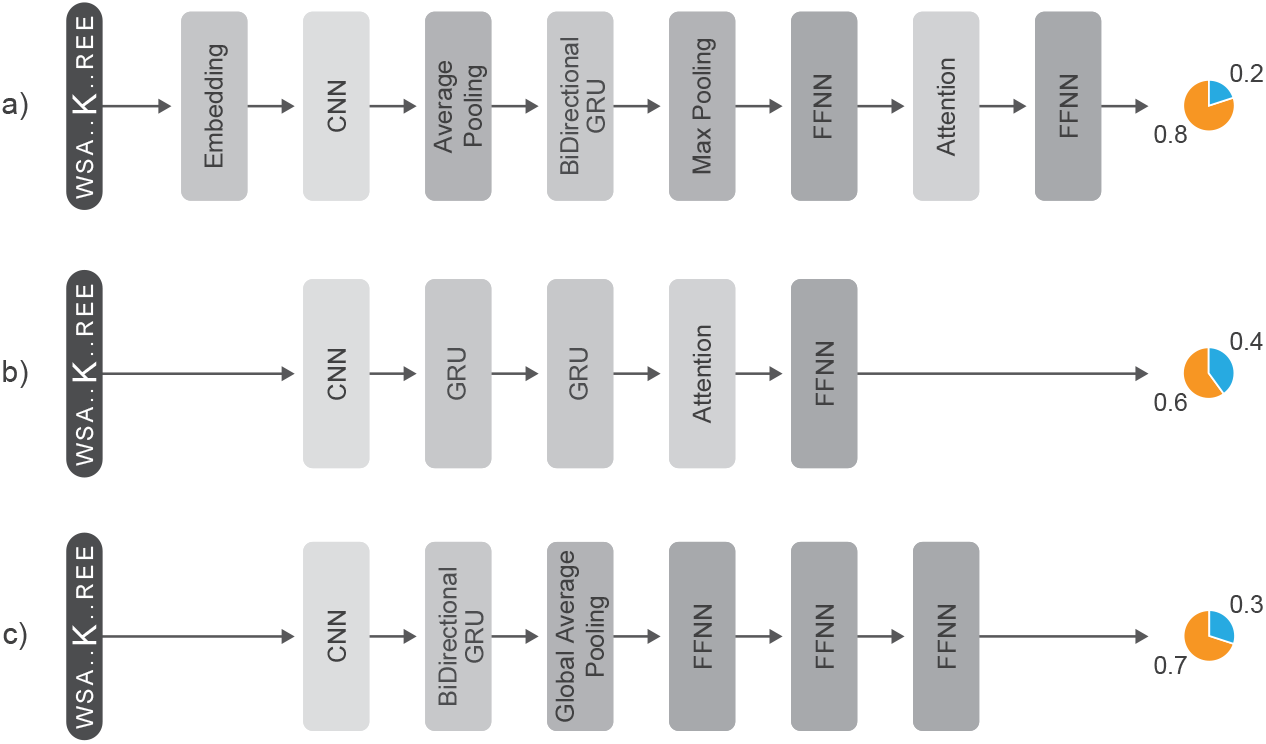
The tree deep learning model architectures : **a)** SUMOnet-1, **b)** SUMOnet-2 and **c)** SUMOnet-3. The input is a peptide sequence centered on the lysine residue, and each model predicts whether the site is going to be predicted SUMOylated (shown in orange) or not (shown in blue).

All CNN layers are 1-D CNNs in all the architectures designed. Pooling layers, average-pooling, maxpooling, and global average pooling are used to decrease model complexity. We apply the ReLu activation function after each CNN and FFNN layer. GRU layers are activated by `tanh` function and we estimate the target class probabilities using `Softmax` function. We optimize the weights of our models with Adam optimizer [28] by minimizing the cross-entropy loss. Below we provide details for each method.

#### SUMOnet-1

SUMOnet-1 performed best when coupled with one-hot encoded vectors as an input. Each sequence is given into a trainable embedding layer to learn better vector representations for input peptides. Then, the CNN layer extracts the information among residues. The extracted features are averaged by using an average pooling layer. The next layer is BiGRU; thus, the network processes feature in forward and backward directions. Max pooling, FFNN, and self-attention layers follow BiGRU. Finally, FFNN connects extracted features to the output layer. The hyperparameters are set after the tuning process. The embedding layer maps each amino acid to a 32-dimensional vector space. 128 filter maps are used in the CNN layer and the size of each filter is 4. The number of memory units of BiGRU is determined as 32 for both forward and backward layers. FFNNs consist of 64 and 256 hidden units, respectively.

#### SUMOnet-2

The best peptide encoding scheme that performed best with the SUMOnet-2 architecture was NLF. Each input vector directly feeds to the CNN layer to extract residue features, which is followed by two GRU layers. The extracted features by CNN are learned sequentially, and the positional information can be captured. Then, the self-attention layer is used for attending to important features by giving weights to each output state of GRU. In the end, FFNN follows self-attention as a final layer. CNN layer consists of 64 filters with a kernel size of 3. We determined 32 units for GRU layers and 128 hidden units for the FFNN layer after the tuning process.

#### SUMOnet-3

SUMOnet-3 performed best when BLOSUM62 encodings are used as input vectors. Inputs are given to the CNN layer to capture features, which is followed by BiGRU. Then, global average pooling is used to calculate the average of feature values and to decrease the number of trainable parameters. Finally, three consecutive FFNN layers are used. After the hyperparameter tuning, the number of filters is determined as 128 with kernel size 2 for the CNN layer. BiGRU consists of 32 memory units, each in forward and backward layers. There are 64, 128, and 128 hidden units in FFNN layers, respectively.

To arrive at these three models, we follow an architecture design process and a hyperparameter tuning schedule, which we detail below.

### 2.4 Architecture Design and Hyperparameter Tuning

The starting point is the shallowest architecture for each encoded input vector. After that, we deepen the network by adding layers that improve the evaluation result one after the other. We use several combinations of well-known neural network units FFNN [3], CNN [29], RNN [39], LSTM [24], GRU [10] and self-attention [50] with pooling methods max-pooling, average-pooling and global versions of them [17]. Further, we add a dropout layer [42] on different parts of the architecture to prevent over-fitting.

We created a search space and tuning schedule for hyperparameter tuning. The tuning schedule consists of searching for the best hyperparameter parameter for a single layer. To do that, the hyperparameters of all layers are set to the value shown in Table 1 except when that specific layer is tuned. The search spaces for each hyperparameter are listed in Table 2. We also tuned the hyperparameters of XGBoost and logistic regression to get the best prediction results. For all models, we used stratified 5-fold cross-validation on the training data and select the best values that maximize the AUC score.

**Table 1:** The neural units that we experimented with and the values that they are set when they are fixed in tuning. - in the cell indicates that there is no parameter to be tuned for that hyperparameter. Adam optimizer uses 0.001 learning rate.

| Neural Units | Fixed Values |
| --- | --- |
| FFNN | 128 |
| 1-D CNN | Filter Size = 120, Kernel Size = 2 |
| RNN | 64 |
| LSTM | 64 |
| GRU | 64 |
| BiDirectional RNN/LSTM/GRU | 64 |
| Self-Attention | - |
| Max-Pooling | 2 |
| Average-Pooling | 2 |
| Global max/average Pooling | - |
| Dropout | 0.3 |
| Optimizer | Adam |
| Batch Size | 32 |
| Epoch Number | 1000 + Early stoppings |

**Table 2:**
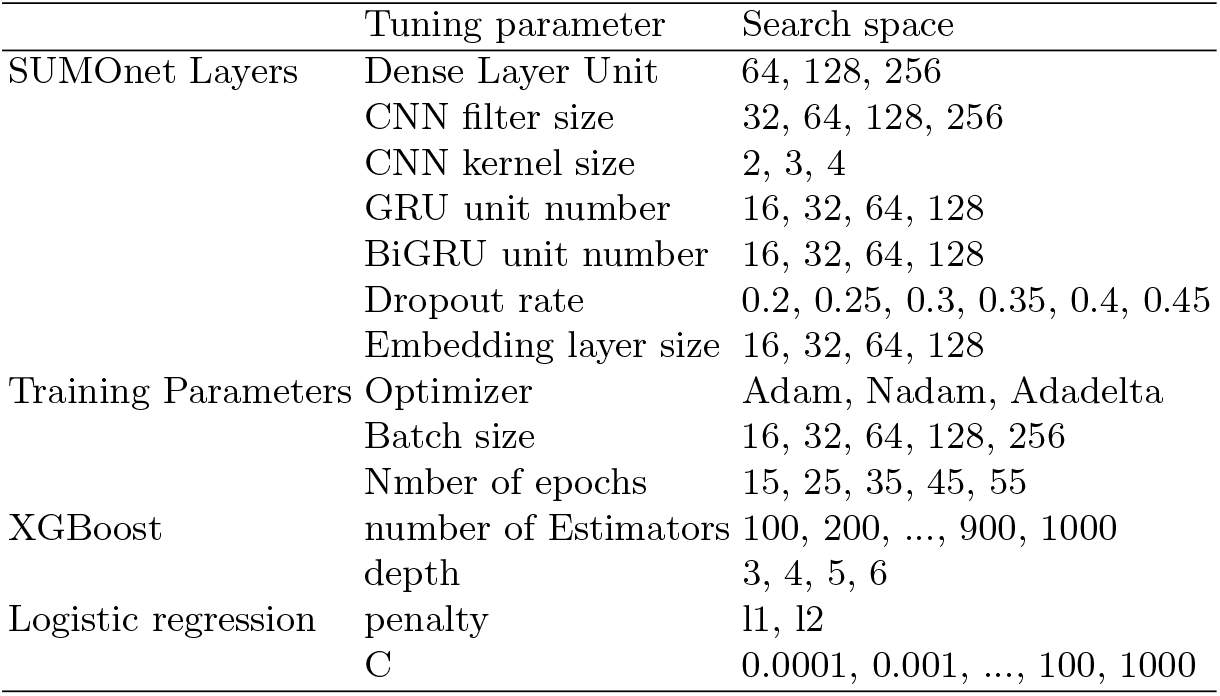
The search space for tuning hyperparameters of each SUMOnet architecture, training components, XGBoost, and logistic regression. We use default learning rates for each optimizer.

## 3 Results

### 3.1 Evaluation Setup

We compare SUMO-nets with three different sets of algorithms:

1. **Rule-based motif classifiers:** To assess what can be achieved by only using the known SUMOylation sequence motifs, we use rule-based classifiers that would predict a SUMOylation site if the SUMO motif is present in the sequence and predict negative class otherwise. We obtain the SUMOylation motifs from [2]. A classifier is built for each motif. We also use an ensemble classifier that would return a positive result if at least one motif matches the input sequence. Since these classifiers will return only binary class labels and no scores to rank instances, we report the evaluation metrics that do not depend on a ranking.
2. **Non-deep learning machine learning classifiers:** To assess if there is any merit in using deep learning models for SUMOylation prediction, we compare SUMOnets with two machine learning methods, logistic regression [35] and gradient boosted decision trees that we build and train. For the latter, we use the XGBoost implementation [5]. We did not make the list of algorithms an extensive one because XGBoost typically outperforms many other algorithms. XGBoost and logistic regression models are also trained with three different encoding methods on the training data. Among the three peptide encodings, both models provide the best performance with BLOSUM62 encoding. Hyperparameters of logistic regression and XGBoost models are tuned on training data using 5-fold cross-validation. The details of the hyperparameter tuning steps are provided in Section 2.4.
3. **Existing SUMOylation predictors:** We compare SUMOnets with the widely available SUMOylation prediction tools as comprehensively as possible. These tools include GPS-SUMO [52], pSumo-CD [27], JASSA [2] and SUMO-Forest [38]. There are many tools developed for SUMOylation prediction and we attempted to include them in our evaluations, but we were hindered by the lack of working implementations or the lack of sufficient details in the descriptions to reproduce these tools independently by ourselves. Some details on how we obtain predictions for each tool we were able to compare are provided below:
  – **SUMO-Forest:** We train SUMOForest on our training data using their available implementation at https://github.com/sandyye666/SUMOForest. The trained model is run on the test data to obtain the predicted class labels and scores.
  – **GPS-SUMO:** The GPS-SUMO [52] source code is not available. Therefore, we use their web server available at [GPS-SUMO web server](http://sumosp.biocuckoo.org/online.php). The test data is uploaded to the webserver to obtain SUMOylation site predictions. The web server provides predicted labels and the associated scores based on a predefined threshold value of the `high, medium, all, low`, and `none`. We apply a medium threshold and use the prediction labels to measure Matthews correlation coefficient (MCC) and F1-Score at this threshold. However, at this threshold, the server does not provide scores for negatively predicted cases. To be able to obtain scores for all instances of ROC and AUPR curves, we use the scores obtained with the threshold setting `all`. The threshold selections are suggested by the authors (via e-mail communication).
  – **JASSA:** The source code is not available; therefore, we used the JASSA webserver at http://www.jassa.fr/. The server takes a single sequence as input. To overcome this issue, we used a Python script to submit test sequences one by one and retrieved each prediction from the results page. Jassa server provides prediction labels and their scores with respect to thresholds `high` and `low` for three different clustering methods, `all, directed` and `inverted`. We used the clustering method `all` and applied both `high` and `low` thresholds separately and calculated MCC and F1-Score. To be able to hold comparisons based on ROC and AUPR curves, we use the scores obtained when applying the clustering method all.
  – **pSumo-CD:** [27] is another widely used SUMOylation predictor. Since the implementation is not available, we resort to the pSUMO-CD server. The server provides prediction labels but not the prediction scores. Thus, for that method, we could only calculate MCC and F1 metrics but not AUC or AUPR-related metrics.

### 3.2 Performance Comparison on All Test Data

We first evaluate each SUMOylation motif performance through rule-based classifiers. These results are presented in Table 3. The maximum F1 score that can be achieved with a single motif is 0.418; the weak consensus motif classifier yields this result. The MCC motif score of this motif is also high, 0.373. The consensus motif achieves 0.403 F1 and 0.374 MCC scores. In general, consensus motifs produce higher scores. Even when all the motifs are considered (the ensemble motif), the highest F1 and MCC scores remain at 0.469 and 0.359, respectively. These results establish that the linear sequence motifs are insufficient to predict SUMOylation sites and form a baseline to compare other methods.

**Table 3:** Evaluation results rule-based motif classifiers on the held-out test set. The motifs are obtained from [2]. The input sequence is predicted as SUMOylated if the corresponding motif matches the sequence. For the ensemble motif, the protein site is predicted as SUMOylated if the protein sequence consists of at least one of the motifs. *Ψ*_1_ = I, L, V; *Ψ*_2_ = A, F, I, L, M, P, V, W; *Ψ*_3_ = A, F, G, I, L, M, P, V, W, Y. *α* = D, E, _*p*_S/T = phosphorylated serine/threonine.

|  | Name | Motif | F1-Score | MCC |
| --- | --- | --- | --- | --- |
| **Consensus direct** | Strong consensus | $[\Psi_1]-[K]-[x]-[\alpha]$ | 0.308 | 0.336 |
| | Consensus | $[\Psi_2]-[K]-[x]-[\alpha]$ | 0.403 | **0.374** |
| | Weak consensus | $[\Psi_3]-[K]-[x]-[\alpha]$ | **0.418** | 0.373 |
| | PDSM | $[\Psi_2]-[K]-[x]-[\alpha]-[x]_2-[S]-[P]$ | 0.015 | 0.076 |
| | NDSM | $[\Psi_2]-[K]-[x]-[\alpha]-[x]-[\alpha]_{2/6}$ | 0.158 | 0.237 |
| | HCSM | $[\Psi_4]_3-[K]-[x]-[E]$ | 0.071 | 0.160 |
| | SC-SUMO | $[P/G]-[x]_{(0-3)}-[I/V]-[K]-[x]-[E]-[x]_{(0-3)}-[P/G]$ | 0.063 | 0.158 |
| | Minimal SC-SUMO | $[I/V]-[K]-[x]-[E]-[x]_{(0-3)}-[P]$ | 0.098 | 0.199 |
| | SUMO-acetyl switch | $[\Psi_2]-[K]-[x]-[\alpha]-[P]$ | 0.092 | 0.180 |
| | pSuM | $[\Psi_2]-[K]-[x]-[pS]-[P]$ | 0.000 | 0.000 |
| **Consensus inverted** | Strong consensus | $[\alpha]-[x]-[K]-[\Psi_1]$ | 0.077 | 0.068 |
| | Consensus | $[\alpha]-[x]-[K]-[\Psi_2]$ | 0.151 | 0.108 |
| | Weak consensus | $[\alpha]-[x]-[K]-[\Psi_3]$ | 0.176 | 0.119 |
| **Ensemble motif** |  |  | **0.469** | **0.359** |

We next evaluated the SUMO-nets with the other two sets of methods, the machine learning models, other deep learning models, and the available SUMOylation predictors. Table 4 shows the evaluation metrics computed on the same test data for all the models. SUMO-Forest is the best predictor among the existing models in the literature. It achieves a 0.59 F1 score, which is approximately 14% higher than JASSA, 17% higher than GPS-SUMO, and 26% higher than pSumo-CD. SUMO-Forest achieves a 0.50 MCC score, which corresponds to 25-30% more than other tools in the literature. As pSumo-CD does not provide scores, some evaluation metrics are missing for this tool; therefore, our interpretation is limited for this method. The ROC-AUC and AUPR scores of SUMO-Forest are 0.82 and 0.69, respectively, which are approximately 11% higher than that of GPS-SUMO’s. Thus, we conclude that SUMO-Forest performs best among the available tools in the literature on these datasets.

**Table 4:** Comparison of the SUMOylation prediction methods on the whole test data. JASSA predictions for two different thresholds are provided, `low` and `high`. pSumo-CD [27] gives the predicted labels without any score; therefore, ROC-AUC and AUPR scores cannot be calculated. Encodings used for the models that we trained are provided in the second column; since other models in the literature do the encoding process themselves, we did not specify them.

| Predictor | Encoding | F1-Score | MCC | AUC | AUPR |
| --- | --- | --- | --- | --- | --- |
| XGBoost | BLOSUM62 | 0.614 | 0.536 | 0.844 | 0.727 |
| LogisticRegression | BLOSUM62 | 0.591 | 0.509 | 0.827 | 0.696 |
| GPS-SUMO | - | 0.428 | 0.157 | 0.707 | 0.569 |
| JASSA-Low | - | 0.403 | 0.380 | 0.727 | 0.560 |
| JASSA-High | - | 0.256 | 0.305 | 0.727 | 0.560 |
| pSumo-CD | - | 0.332 | 0.215 | - | - |
| SUMO-Forest | - | 0.592 | 0.502 | 0.819 | 0.688 |
| SUMOnet-1 | One-Hot | 0.640 | 0.544 | 0.859 | 0.742 |
| SUMOnet-2 | NLF | 0.623 | 0.547 | 0.857 | 0.741 |
| SUMOnet-3 | BLOSUM62 | **0.658** | **0.569** | **0.870** | **0.758** |

We trained logistic regression and XGBoost models to set a strong baseline for the deep learning models. Our results show that the methods in the literature fall behind these two models in all metrics. XGBoost produces about a 2% improvement in performance for all the evaluation metrics over logistic regression. For the XGBoost classifier, the F1-Score and ROC-AUC are 0.61 and 0.84, respectively, which are 2% higher than SUMO-Forest. The MCC and AUPR scores are 0.54 and 0.72, respectively. This corresponds to a 3% increase compared to the best tool in the previous comparison, SUMO-Forest. Thus, we conclude that a SUMOylation predictor better than the existing models can be trained with an XGBoost classifier.

Finally, we examine SUMOnets prediction performance Table 4. All three SUMOnets outperform the models in the literature and the trained XGBoost model. Among the three architectures, SUMOnet-3 is the one that achieves the best scores. The F1-Score of SUMOnet-3 is 0.66, which is 5% more than XGBoost classifier [5]. MCC, ROC-AUC and AUPR scores of SUMOnet-3 are 0.57, 0.87 and 0.76, respectively. These correspond to an improvement of approximately 3% over the XGBoost classifier. We conclude that we can attain the best predictor by using SUMOnet-3. We also compare all the models using the Receiver operating characteristic (ROC) and precision-recall (PR) curves. As shown in Figure 3a SUMOnet-3 is the best overall predictor across different false positive rates.

**Fig. 3:**
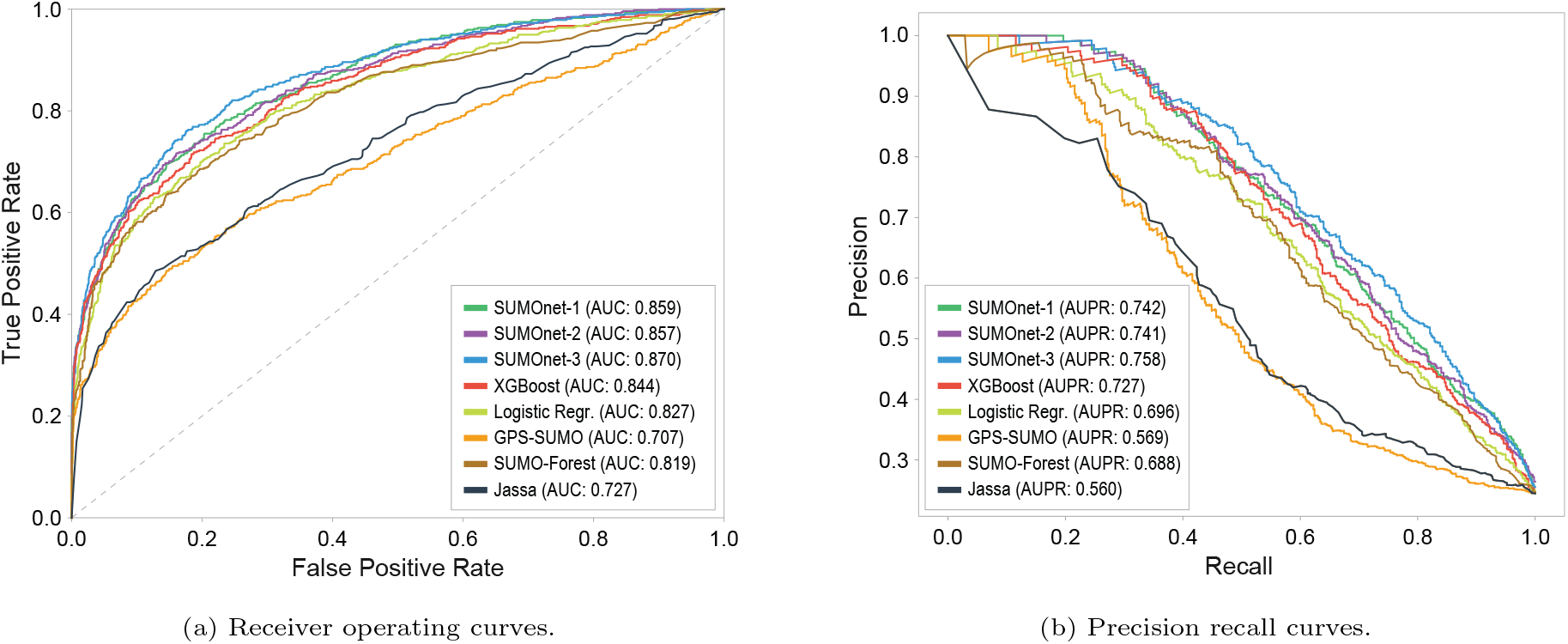
Performance comparison of different models evaluated on the entire test data.

### 3.3 Performance Comparison on Hard Test Examples

The hard test set includes target sequences that lack SUMO motifs but are SUMOylated and sequences that are not SUMOylated but contain a SUMOylation motif. These are the specific examples that would challenge predictors. Table 5 lists the performance scores. The predictive performance of all methods degrade in general. Especially, the methods that rely heavily on the motifs, such as JASSA and GPS-SUMO, lead to predictions worse than a random choice predictor. Figure 4 shows the comparison of ROC curves for various methods. SUMOnet models perform the best compared to both XGBoost and previous methods. SUMOnet-3, which relies on CNN and biGRU, is the best performer among all. This result demonstrates that the deep learning model can go beyond learning simple linear motifs.

**Table 5:** Comparison of the SUMOylation prediction methods on the hard test examples. Hard test examples are the subset of our test data set. We select positive samples which have no motifs and negative samples which have at least one motif.

| Predictor | Encoding | F1-Score | MCC | AUC | AUPR |
| --- | --- | --- | --- | --- | --- |
| XGBoost | BLOSUM62 | 0.438 | 0.0838 | 0.523 | 0.742 |
| LogisticRegression | BLOSUM62 | 0.410 | 0.0598 | 0.507 | 0.740 |
| GPS-SUMO | - | 0.357 | -0.491 | 0.321 | 0.605 |
| JASSA-Low | - | 0.028 | -0.364 | 0.240 | 0.564 |
| JASSA-High | - | 0.000 | -0.259 | 0.240 | 0.564 |
| pSumo-CD | - | 0.194 | -0.091 | - | - |
| SUMO-Forest | - | 0.460 | 0.101 | 0.533 | 0.762 |
| SUMOnet-1 | One-Hot | 0.547 | 0.152 | 0.657 | **0.822** |
| SUMOnet-2 | NLF | 0.479 | 0.118 | 0.601 | 0.787 |
| SUMOnet-3 | BLOSUM62 | **0.565** | **0.168** | **0.667** | 0.819 |

**Fig. 4:**
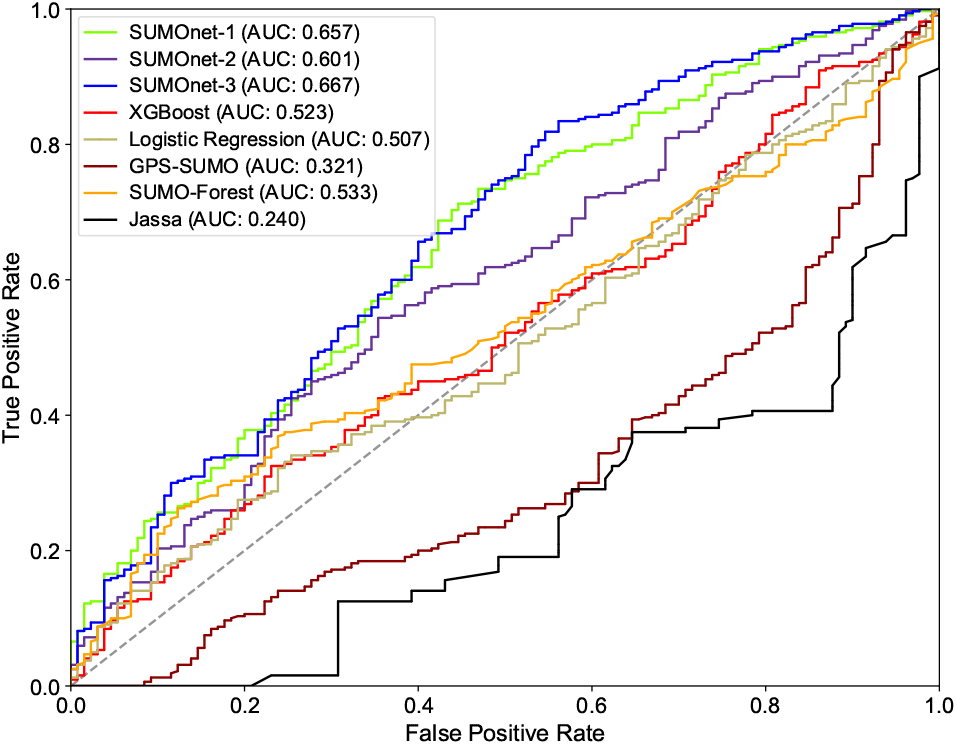
ROC curves of different models on hard test examples. Hard test samples include sequences that lack any SUMOylation motifs but are known to be SUMOylated and sequences that are not SUMOylated but contain a SUMOylation motif.

### 3.4 Ablation Study on SUMOnet-3

To assess the contribution of each component in SUMOnet-3, we add the SUMOnet-3 components gradually and re-assess the model performance. Each model is evaluated using 5-fold cross-validation on the training data. The starting point for the architecture construction is a single layer perceptron that contains 2 hidden units. When the convolution layer is added, we observe an improvement in all the evaluation metrics Table 6. The largest performance increase is observed when BiGRU is added after the CNN, with nearly 3% in ROC-AUC and 4% in AUPR. There is a small improvement in all the scores after adding global average pooling and the three consecutive dense layers.

**Table 6:** Evaluation results of each model as we add components of SUMOnet-3 in the architecture. The table reports 5-fold cross-validation average scores on the training data. The BLOSUM62 encodings are used for input representation.

| Method | F1-Score | MCC | AUC | AUPR |
| --- | --- | --- | --- | --- |
| Dense[2] | 0.538 | 0.492 | 0.831 | 0.697 |
| Conv[120,2]_Dense[2] | 0.584 | 0.507 | 0.837 | 0.709 |
| Conv[120,2]_BiGRU[32]_Dense[2] | 0.633 | 0.545 | 0.864 | 0.745 |
| Conv[120,2]_BiGRU[32]_GlobalAv.Pool_Dense[2] | 0.628 | 0.551 | 0.868 | 0.751 |
| Conv[120,2]_BiGRU[32]_GlobalAv.Pool_Dense[128,128,128,2] | **0.640** | **0.557** | **0.873** | **0.756** |

## 4 Conclusion and Future Work

We present three deep learning architectures, SUMOnet-1, -2, and -3, to predict SUMOylation sites accurately. SUMOnets surpass the closest best SUMOylation predictor available in the literature by a large margin in all evaluation metrics. SUMOnets are built by various neural units that are known to perform well in sequential learning tasks, such as CNN, GRU, and attention. Further, we couple these architectures with different input representations. The proposed architectures provide the most accurate deep-learning architectures for these encoding techniques.

In this work, we also assessed the motif-based rule classifiers that rely solely on the absence and presence of SUMOylation motifs. The results demonstrate that using motifs alone is very limited for accurate prediction of SUMOylation sites. We also train XGBoost and logistic regression models to form a strong baseline. We show that it is possible to attain a good SUMOylation predictor with an XGBoost classifier that outperforms the SUMOylation predictors we compare. We finally observe that SUMOnets that are based on deep sequential architectures achieve better performance among all. We show that among the three SUMOnet architectures, SUMO-net3 achieves the best performance across all metrics. When the methods are compared on a challenging subset of the test data, the performance improvement over the existing SUMOylation predictors becomes more evident. We also observe that widely used predictors that rely on motifs would perform worse than a random choice classifier for such inputs. SUMOnet-3 is the best SUMOylation site prediction tool in all the evaluation experiments. We provide SUMOnet-3 as an open-source project in GitHub and a Python library that can be easily installed by `pypi`. The library also offers means to reproduce our results for further evaluation.

The work can be extended in future directions. Our preliminary experiments with protein sequence embedding methods with ProtVec[1] did not outperform the existing embedding methods we used (data not shown). However, there has been great progress in protein language models [11]. It would be interesting to explore whether alternative embeddings can further improve prediction performance. Lastly, the PTM classification tasks are similar. It could be useful for site prediction for other PTM types. We plan to experiment with these architectures for other PTMs.

## 5 Acknowledgement

We thank Dr. Umut Sahin (Bosphorus University) for inspiring us to work on SUMOylation.

